# Automated Feature Extraction from Large Cardiac Electrophysiological Data Sets

**DOI:** 10.1101/2020.10.21.340968

**Authors:** John Jurkiewicz, Stacie Kroboth, Viviana Zlochiver, Peter Hinow

## Abstract

**Rationale:** A new multi-electrode array-based application for the long-term recording of action potentials from electrogenic cells makes possible exciting cardiac electrophysiology studies in health and disease. With hundreds of simultaneous electrode recordings being acquired over a period of days, the main challenge becomes achieving reliable signal identification and quantification.

**Objective:** We set out to develop an algorithm capable of automatically extracting regions of high-quality action potentials from terabyte size experimental results and to map the trains of action potentials into a low-dimensional feature space for analysis.

**Methods and Results:** Our automatic segmentation algorithm finds regions of acceptable action potentials in large data sets of electrophysiological readings. We use spectral methods and support vector machines to classify our readings and to extract relevant features. We are able to show that action potentials from the same cell site can be recorded over days without detrimental effects to the cell membrane. The variability between measurements 24 h apart is comparable to the natural variability of the features at a single time point.

**Conclusions:** Our work contributes towards a non-invasive approach for cardiomyocyte functional maturation, as well as developmental, pathological and pharmacological studies. As the human-derived cardiac model tissue has the genetic makeup of its donor, a powerful tool for individual drug toxicity screening emerges.

## 1 Introduction

Cardiac function is driven by electrical fluxes that trigger mechanical forces in living tissue. Voltage-gated ion channels regulate the flux of multiple ion species (K^+^, Na^+^, Ca^2+^) that result in subsequent mechanical contraction. Due to this close connection between electrical and mechanical behavior in the heart, the study of the electrical impulses can give valuable insight into the contractile behavior of cardiac tissue, as well as the emergence of arrhythmias (Anumonwo and Pandit, 2015; Qu et al., 2018). The propagating transmembrane potential is known as action potential (AP), and is critically important in the governance of proper cardiac behavior.

Direct recording of APs has a rich and storied history since the foundations of the field of electrophysiology (Hodgkin and Huxley, 1952; Harrington and Johnson, 1973). Due to the need for intracellular access this has proven to be time-consuming and technical, e.g. using the standard voltage-clamp technique (Fozzard and Beeler, Jr., 1975). Such techniques also do not allow for detailed investigation of the connection between transmembrane and extracellular potentials. AP waves in cardiac tissue can be measured indirectly, by means of recording an ancillary potential known as field potential (FP). Much like a medical electrocardiogram, FP recordings measure the potential change across the entire bulk of tissue and have been observed to be closely correlated to the underlying AP (Tertoolen et al., 2018). Recent work has shown the feasibility of multi-electrode arrays (MEA) (Meyer et al., 2012) in the large-scale efficient recording of both AP and FP readings (Edwards et al., 2018; Zlochiver et al., 2019; Hayes et al., 2019; Balafkan et al., 2020).

In a MEA, cardiac tissue is plated onto a substrate situated atop an array of electrodes embedded in the base of a well. By measuring the potential difference between these base electrodes and reference electrodes surrounding the outer perimeter of the well, FPs can be readily recorded in large volumes (Kujala et al., 2011). Further, delivery of electroporating pulses via these electrodes creates pores in the cell membranes and allows direct AP measurements. These readings can be obtained without complicated measuring techniques and provide data from live beating tissue. Edwards et al. (2018) have shown that the myocytes plated on an MEA remain alive and electrically active with consistency after electroporation, in contrast to the terminal effects of microelectrode impalement on cell vitality. This enables testing of long-term effects of different compounds or environments on cell behavior. To the best of our knowledge, a rigorous statistical analysis and comparison of the signals obtained via repeated MEA measurement at different times is currently lacking.

A determining factor in the applicability and efficiency of AP-related assays is the availability of ethically-sourced cardiac tissue for experimentation. The historical precedent in electrophysiology has been the use of animal cells cultured from non-human mammals (Mathur et al., 2016). Nonetheless, differences between nonhuman models and humans could contribute to clinical trial failures despite ideal nonhuman results (Li et al., 2020). The pioneering work of Yamanaka (Takahashi and Yamanaka, 2006) in the reprogramming of adult cells into stem cells has opened a new avenue of cardiac research by allowing the study of human cardiac tissue With this approach, large cell cultures of human tissue can be made available for research purposes. Recent work has shown potential for human induced pluripotent stem cell-derived cardiomyocytes (hiPSC-CMs) (Zwi et al., 2009) to allow for drug testing and disease modeling on viable human cardiac tissue in a controlled environment (Braam et al., 2010; Tertoolen et al., 2018; Edwards et al., 2018; Gintant et al., 2019; Kussauer et al., 2019). The genetic makeup of hiPSC-CMs is identical to that of their donor which makes them a tool to test personalized drug responses (Strauss and Blinova, 2017). While there are concerns regarding the level of maturity of these hiPSC-CMs ranging from fetal to neonatal in development (Zhu et al., 2017), the possibility for safe human-tissue drug testing demands further research.

AP readings are analyzed by encoding the data stream as sequences of features (or biomarkers) that characterize the beat-by-beat behavior of the cells. These features as well as their counterparts in frequency space encode trends in the behavior of the cells. APs are typically encoded as action potential duration at *m* % (APD_*m*_), the time from peak voltage until the cell has repolarized to *m* % of its resting potential. Additionally, differences and ratios of these features are commonly used (Britton et al., 2017). Using these features, the AP data can be projected into a low-dimensional feature space while still retaining the most relevant information for clinical purposes (Cantwell et al., 2019).

Both FP and AP readings are fundamentally time-periodic quantities and it is therefore natural to analyze these signals in the frequency domain. The Discrete Fourier Transform (DFT) (Oppenheim and Schafer, 1999) enables the decomposition of a temporal signal into its constituent frequency components. Analyzing the behavior of these frequency spectra can give indications of signal quality and variance from a defined standard behavior.

The parallel configuration of an MEA allows to generate vast amounts of data in short time. However, this is also one of the challenges of this approach. Regardless of he experimental setup, access to AP data involves delivery of stimulating electrical impulses to the tissue, resulting in large regions of data without electrophysiological meaning. Further, electroporation does not offer permanent intracellular access, as the membrane nanopores close in time. This results in a degradation of the signal as the AP trace reverts to an FP morphology. Electroporation may even fail entirely for a given cell and yield no usable AP data. The large volume of data generated thus requires a robust segmentation protocol to extract usable data. Manual segmentation is time-consuming and error prone for even modest data sets and becomes quickly infeasible as the volume of data grows. Therefore, the need for an automated segmentation method is imminent.

Here we propose an automated paradigm to efficiently analyze large sets of MEA data, perform segmentation and extract features. The method relies on analysis of the time derivative of the AP. A graphical depiction of the workflow is found in Figure 2. We further utilize this paradigm to extract features from a multitude of hiPSC-CMs for purposes of statistical analysis. We apply the algorithm to extract data from the same cell site at different time-stamps to analyze the similarity for repeated physiological measurements. Data loading occurs prior to implementation, and the open source nature of the algorithm ensures that the program can be readily adapted to any specific setting and format of input data. Thus, the method is capable of determining signal quality for virtually any type of experimental regimen for data collection, not exclusively MEA experimentation.

**Figure 1:**
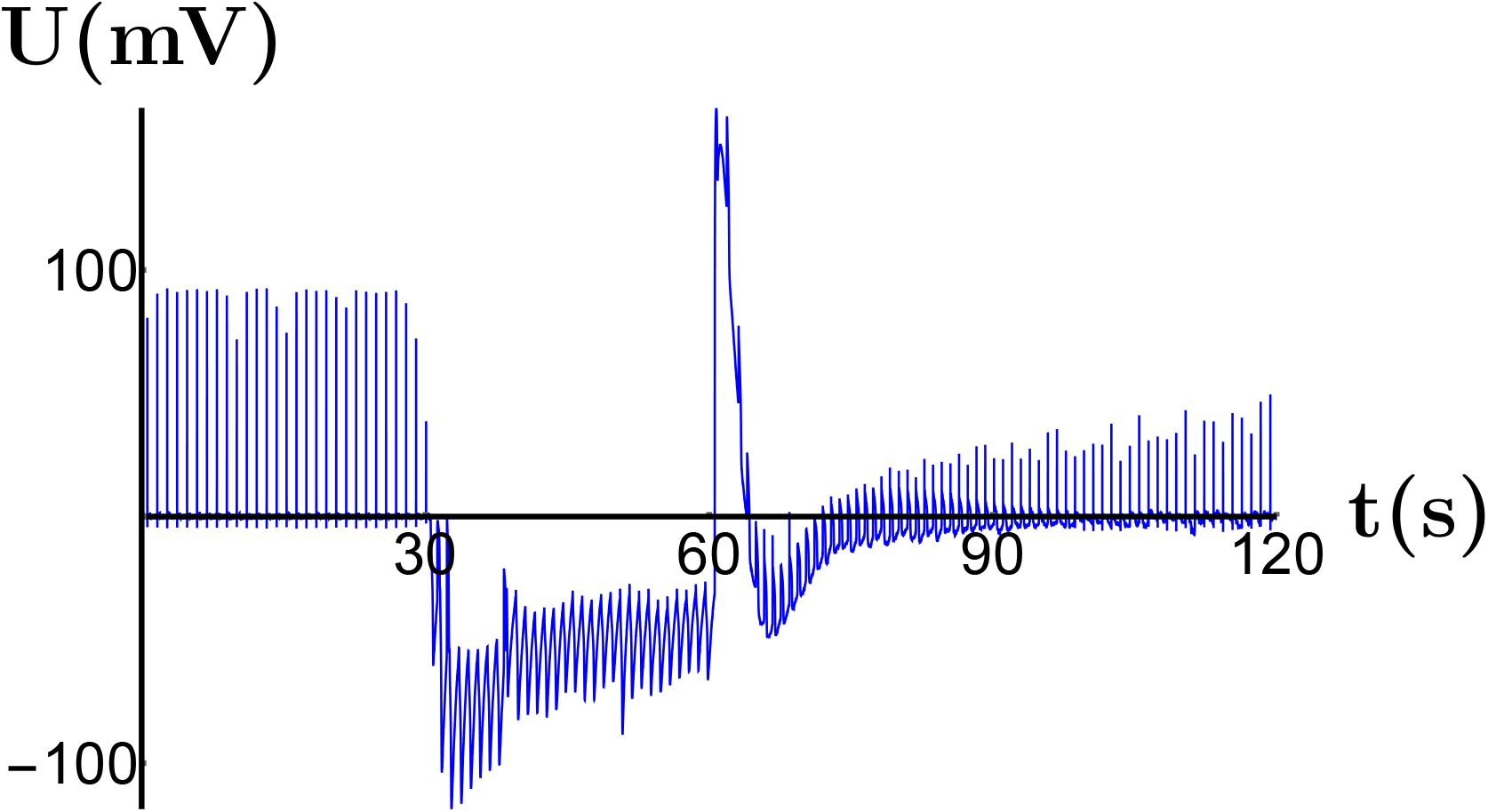
Raw data stream from a single electrode in an MEA. Prior to 30 s, the signal consists exclusively of FPs. From 30 s to 60 s, the electroporation stimulus is applied. From 60 s to 120 s, the signal includes APs that gradually degrade back to FP morphology.

**Figure 2:**
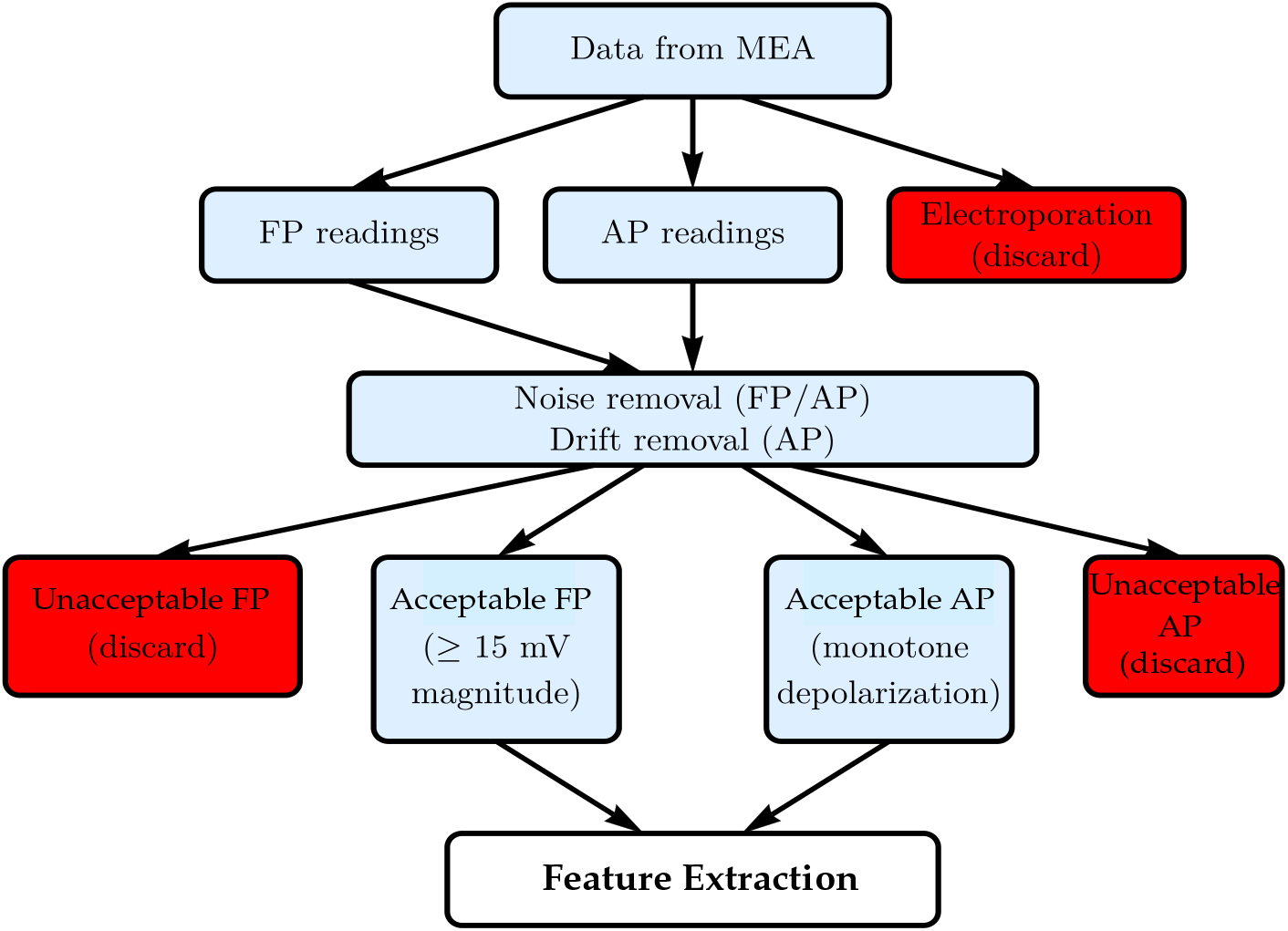
A flow chart showing the workflow for the segmentation algorithm.

## 2 Materials and Methods

### 2.1 Experimental Setup

High-purity iPSC-CM cultures were produced from human cardiac fibroblasts as described in (Zlochiver et al., 2019). hiPSCs cultures were differentiated using the feeder-free monolayer differentiation protocol. RPMI with B27 minus insulin containing 6 *µ*M CHIR99021 (4423, R&D Systems) was added to the cultures at time zero. After 3 days, the cultures were treated with 5 *µ*M IWR-1 (I0161, Sigma Aldrich) for 48 h. Subsequently, the cultures were maintained in RPMI media with B27 without insulin for 5 days and then with RPMI with B27 complete supplement for cell maturation.

PEDOT-coated 24-well micro-gold multielectrode array (*µ*GMEA) dishes (catalog number 24W300/30G-288, MultiChannel Systems MCS, Germany) with an array of 12 electrodes per well underwent a 30-min hydrophilic treatment with fetal bovine serum (FBS) and fibronectin-coating (50 *µ*g/mL) for 3 hr at 37 ^°^C prior to cell plating. hiPSC-CMs cultures were then dissociated using Accutase (Sigma) enzyme and plated at 30,000 cells/5 *µ*L/*µ*GMEA well. Spontaneous beating of cardiomyocytes was first observed in 2-day-old constructs, see the video in the Supplementary Material.

FPs and APs were recorded daily from 2 to 15-day-old constructs and APs from 10-day-old constructs at a sampling rate of 20 kHz. For experiments where electrodes were stimulated, an electroporating pulse (1 V, 1 Hz, and 1 ms) was delivered for 30 s. The raw data are available from the corresponding author upon request.

### 2.2 Data Segmentation and Feature Extraction

The purpose of this algorithm is to identify a given signal stream as AP or FP, determine acceptability of the signal and to extract the maximum amount of acceptable AP signals. Unacceptable APs feature a non-monotone depolarization due to partial closure of the nanopores, see Figure 3 (A). When the data are obtained via MEA experimentation, additional FP data are extracted. In each experiment the time of initiating electroporation and the duration of the pulse are recorded. Hence the region prior to the electroporation timestamp consists of FP readings, while the region following cessation of the pulse contains AP readings of varying quality. From these two regions, candidate traces of FP and AP are extracted. Acceptable FPs are those that exceed a threshold of 15 mV.

**Figure 3:**
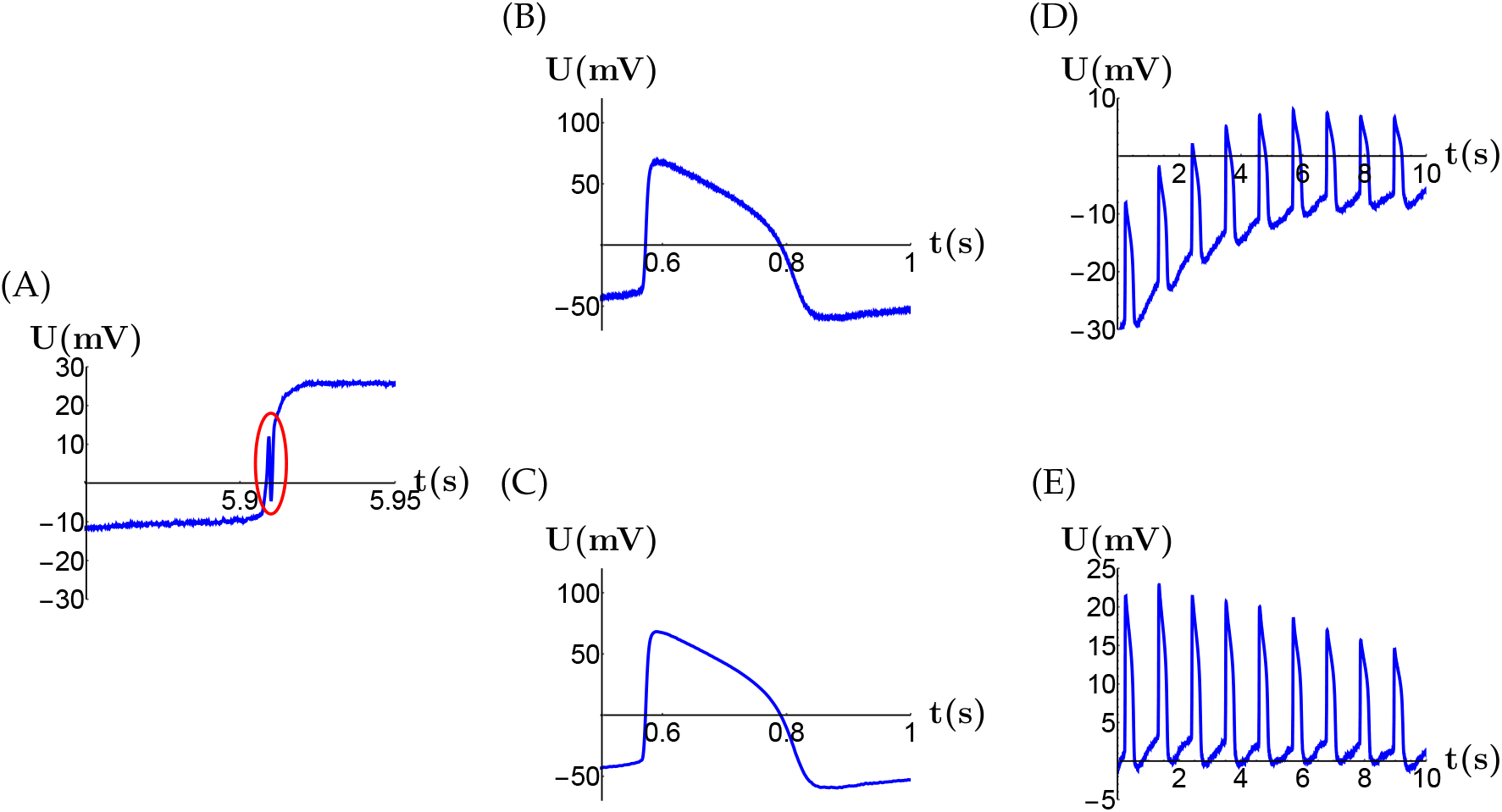
(A) Unacceptable AP trace featuring non-monotone depolarization. (B) & (C) AP trace before and after application of digital filter. (D) & (E) AP trace before and after removal of baseline trend.

Signal classification is accomplished in the Fourier space. Let **U** denote the voltage signal, **Û** its DFT, and **Û** _**200**_ be the portion of the DFT with argument below 200 Hz. We encode **Û** by its *L*^2^-mass (or energy) and the fraction of the mass below 200 Hz, i.e.

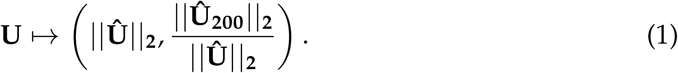

We use a neural network to classify signals under this encoding. We formulate a training set by encoding 494 signals - 255 FP and 235 AP - which demonstrate that these signals are linearly separable. That is, there exists a line *ax* + *by* + *c* = 0 such that all FPs are located on one side and all APs are located on the other side of that line. We choose a support vector machine (SVM) network to compute the line of maximal margin (Fine, 1999). Our SVM is trained on this data via quadratic programming (Decoste and Schölkopf, 2002). Due to linear separability, the classifier is 100% accurate on the training set (Fine, 1999).

Electrical noise and baseline trends in the APs are removed prior to determining the data quality, see Figure 3 (B)-(E). We use two Butterworth filters (Butterworth, 1930) applied sequentially, a low-pass filter with cutoff frequency 225 Hz for noise removal and a high-pass filter with cutoff frequency of 0.5 Hz for removal of baseline trend. Each filter is of a different order to maintain stability and optimize computational performance. The processed data are numerically differentiated using centered differences. The algorithm locates positive maximums of the derivative due to repolarization and possible significant negative minimums in a neighborhood of a local maximum. The signal is truncated at the earliest instance of non-monotonicity. The algorithm is written in the Open Source language Python (Python Software Foundation, 2020), carefully documented and available at https://github.com/jurkiew4/Action-Potential-Segmentation.

Once the signal has been processed, acceptable data streams are submitted for feature extraction. We compute three classical features of APs, namely basic cycle length (BCL), APD_30_ and APD_80_. The BCL is computed using the same program that locates depolarization spikes from the segmentation step, followed by finding the time between successive maximums of the derivative. The APD_*n*_ is computed by isolating each individual pulse as above. Each pulse is shifted and re-scaled so that the maximum voltage occurs at time zero with magnitude 1, while the minimal value of the voltage for that pulse is shifted to a value of 0. Frequency spectra are extracted by taking a 15 s segment of each signal starting at *t* = 0 s for that signal and computing the DFT for that window. The modulus of the complex frequency component is then plotted against the frequency. The 15 s window moves in steps of 0.5 s.

## 3 Results

Running the segmentation program on 864 candidate data streams from untreated cells (3 full MEA plates of 288 electrodes each) yields 408 coupled pairs of acceptable processed AP and FP traces. The duration of the segmentation per plate is a few minutes on a commercially available personal laptop, running an Intel i7 processor with 8 GB RAM. The AP traces vary in length from 5 to 50 s. Representative coupled 15 s traces are shown in Figure 4.

**Figure 4:**
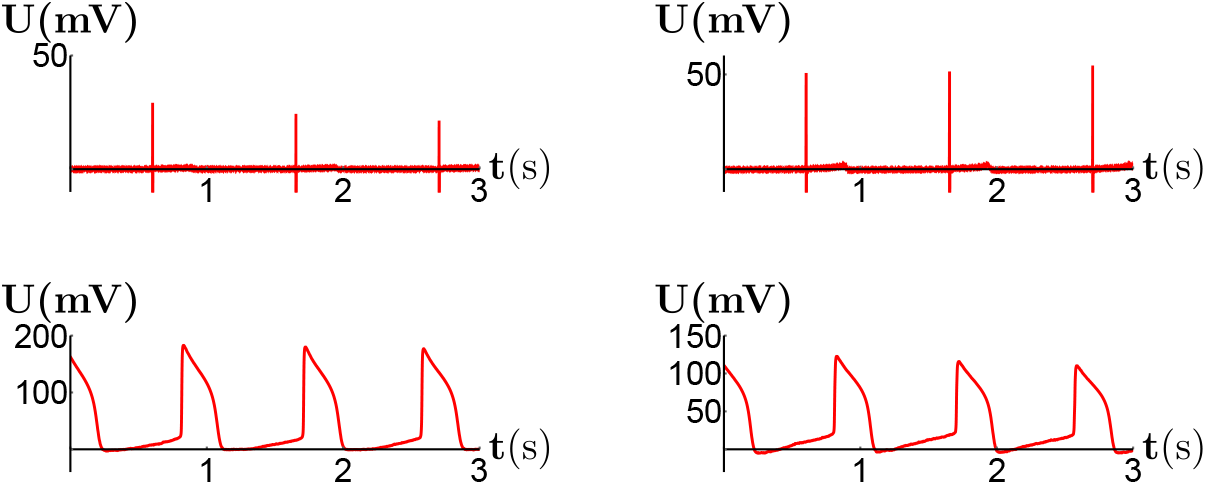
Coupled FP (top) and AP (bottom) readings from the same electrodes. Note the slightly higher beat rate of AP vs. FP, a result of the 1 Hz electroporation stimulus.

The dynamic behavior of the features as time series for each electrode manifest the signal degradation to an FP-like trace and a return of the BCL to an intrinsic beat rate in the absence of electroporation stimulus (5). Reversion to FP-like behavior sets in after between 4 and 50 beats, while on average, the nanopores remain open for about 30 s. In the longer time series we observe increasing BCLs and increasing APDs, indicating that the intrinsic beat rate of these cells is less than 1 Hz. Remarkably, even large fluctuations in BCL frequently do not manifest similar fluctuations in APD. This implies a largely uniform AP morphology in spontaneously beating cells.

A two-dimensional encoding of the frequency spectrum of the APs and FPs allows a clear distinction between them. Figure 6 shows the separation that these features create for 494 traces, 255 FP traces and 235 AP traces. Of the AP traces, 175 are from MEA readings from hiPSC-CMs and 60 are *in silico* AP traces generated using the Grandi et al. (2010) model with randomized pacing protocols. Note that the margin is actually a conservative choice as it is determined by two isolated data points at the upper end of the FP cloud.

**Figure 5:**
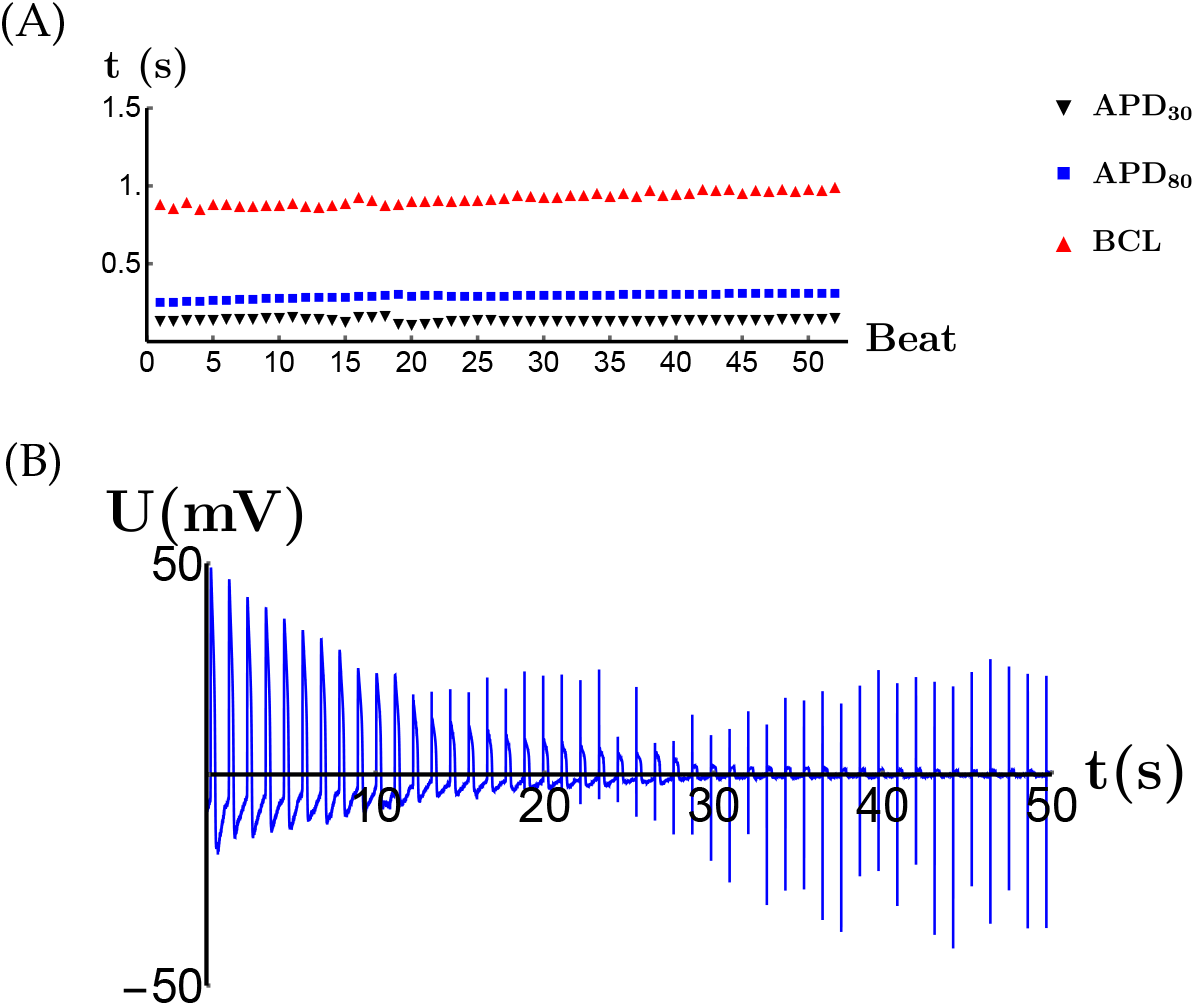
(A) Time series plot of APD_30_, APD_80_ and BCL for a single data stream. (B) An AP signal chain of 50 s recorded after electroporation on an MEA. The unstable nanopores begin closing after approximately 13 s, resulting in a transitional signal morphology. After 30 s the nanopores are almost fully closed and the signal has reverted to an FP-like morphology.

**Figure 6:**
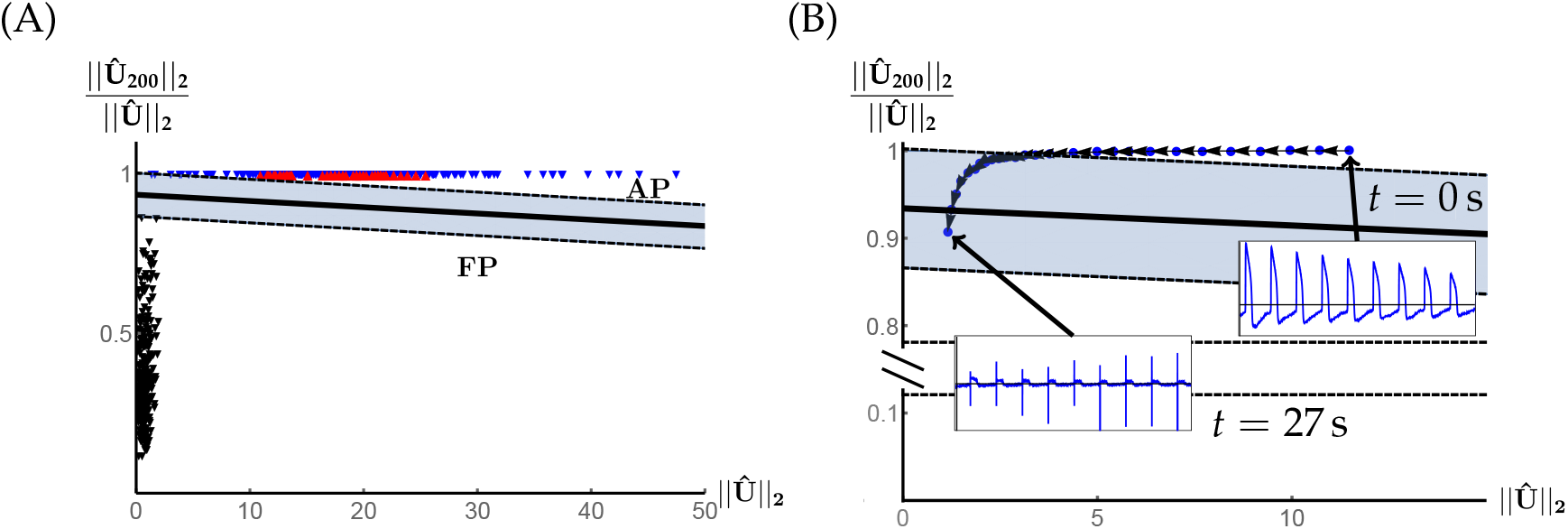
(A) The two frequency features from Equation (1). This choice allows linear classification of a signal morphology as a sequence of FPs or APs. In vitro AP recordings are marked by blue triangles, and *in silico* simulations are marked by red triangles. The bold line indicates the line of maximal margin obtained via SVM training. The region between the maximal margins on either side is shaded in gray. A signal with Fourier features in this region is discarded as neither AP nor FP. (B) Feature-space encoding of signal train in Figure 5(B). Each successive state represents an increment of ∆*t* = 1 *s* in the signal. The two-dimensional encoding allows for a refined view of the signal degradation to FP-like behavior.

In Figure 7(A) we show AP traces from the same electrode recorded 25 h apart. To analyze similarity and reproducibility of measurements across days, we run the segmentation algorithm on data sets from the same MEA plate at 3 different times, *t* = 0, 48 and 96 h, respectively. Figure 7(B)&(C) shows a strong coincidence of the APD_80_ and BCL distributions. Treating each sequence of APDs from each day as a random sample from a probability distribution, we can form the empirical density function for each distribution. Measuring the pairwise distances between the empirical distributions on each day in the supremum norm we observe a maximal distance of 0.3, similar to the maximal supremum distance between distributions of different cell batches. Zhu et al. (2016) have shown that populations of stem cell-derived myocytes can exhibit significant variability in mean APD, and our results conform to these observations. Moreover, any day-to-day variance in APD readings is comparable to the natural variability of readings at a single time point.

**Figure 7:**
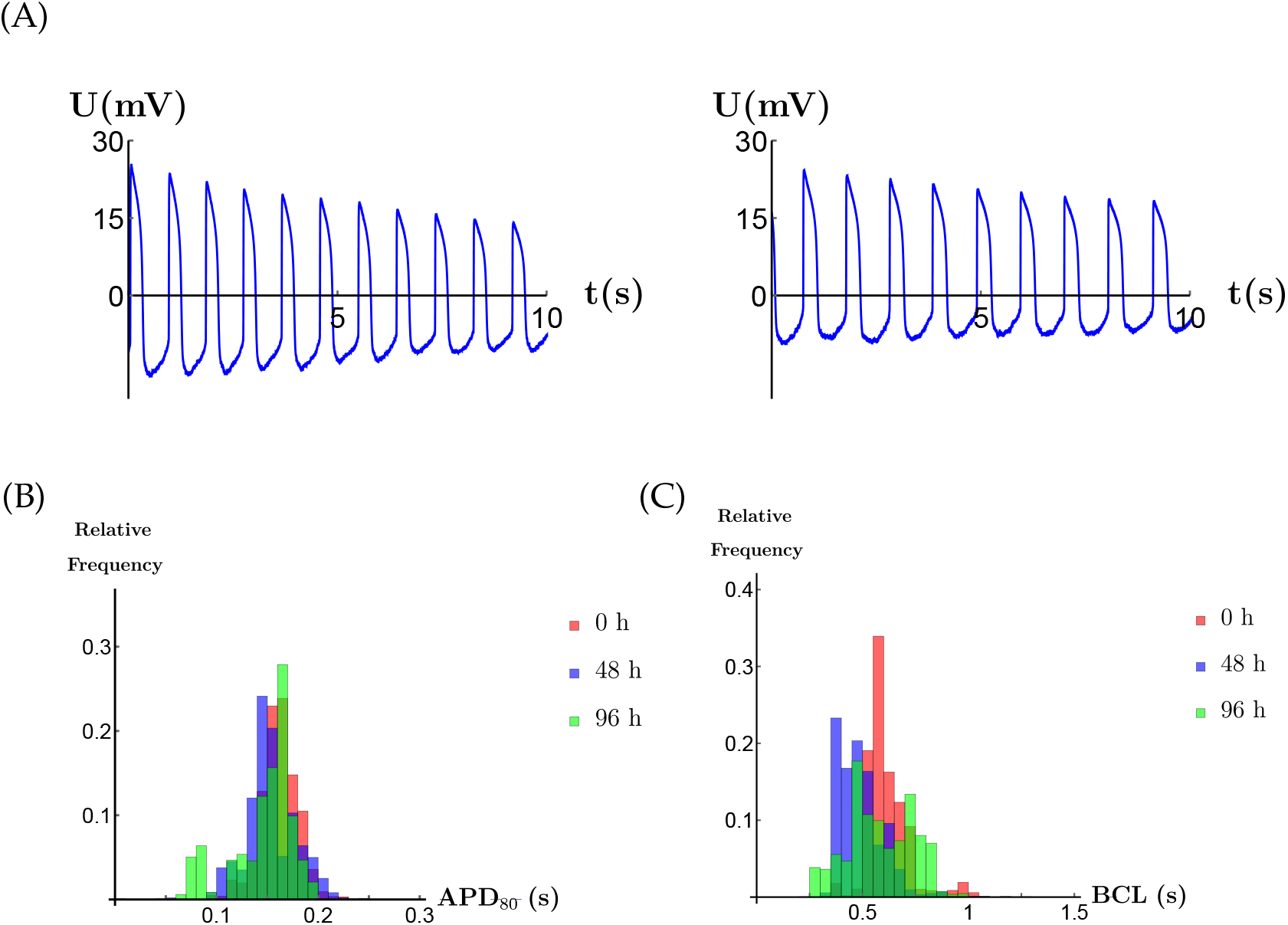
(A) AP traces from the same cell site taken 25 h apart, both recorded 10 s postelectroporation. The similarity in morphology and magnitude indicates consistency of measurements across multiple experiments. (B) & (C) Distributions of APD80 and BCL collected from the same MEA plates at 0, 48 and 96 h, respectively.

## 4 Clinical Relevance

A new paradigm based on hiPSC-CMs has been proposed for personalized prediction of proarrhythmic risk (Li et al., 2020). As hiPSC-CMs retain the genetic makeup of their donor, they provide a powerful tool for drug toxicity screening, understanding individual disease phenotypes and developing precise therapeutic strategies.

Experimentation with hiPSC-CMs models has identified ion-trafficking defects associated with pathological mechanisms of cardiovascular disease. Ion channelopathies are one of the most well-established iPSC-based disease models because of their well-understood impact on the AP (Fernández-Falgueras et al., 2017). Abnormalities occurring in AP generation, synchronization, or propagation may cause cardiac channelopathies related to arrhythmia (Smith et al., 2017). Different disease-specific human iPSC lines have been developed from patients with ion channel gene mutations (Takaki et al., 2019; Wuriyanghai et al., 2018; Malan et al., 2016; Park et al., 2019). Given that the regulation of key genes may be governed by a complex regulatory landscape, relevant human cell disease models are of vital importance for precision medicine. Our algorithm provides the field of translational cardiac electrophysiology with a robust and efficient tool to identify and extract usable data from the large recording volumes necessary to obtain statistically reliable information.

## 5 Discussion and Conclusion

MEA technology for industrial-scale recording of coupled FP and AP data allows for sophisticated and extensive analysis of electrophysiological phenomena. Combined with the introduction of induced pluripotent stem cells by Yamanaka in 2006 (Takahashi and Yamanaka, 2006), this creates the potential for vast amounts of data to be collected in a short period of time.

To parse the significant amount of data generated by even a single MEA experiment and determine which readings are useful sound for analysis, we examine the signal in a 2-dimensional feature space and inspect the time derivative of the signal. Our algorithm efficiently locates and classifies AP and FP signals and determines the acceptability of the signal using only consumer-grade hardware. This allows for quick extraction of relevant data from an experiment without human intervention, freeing up time for experimental analysis. Our algorithm also efficiently addresses the random duration of intracellular access and the slow reversion of AP signals to FP morphology. This enables maximal information extraction for a given experiment. By analyzing the derivative to locate the onset of degradation, we eliminate dependence of the algorithm on a specific experimental procedure and make it applicable to non-electroporation based MEA methods, such as those proposed by Hayes et al. (2019).

MEA technology allows for repeated large-scale electrophysiological readings on the same cells at different timestamps, enabling detailed analysis of the effect of time on various physiological readings. We observe that the variability in readings taken from the same cell site at different days is comparable to the variation in different cell populations and thus is not an impediment to the reproducibility of experiments over timescales on the order of days. This long-scale consistency represents a significant strength of the MEA approach to electrophysiology and its applicability to clinical research.

The large amount of data that can be extracted from this experimental setup also allows for detailed analysis of the behavior between features for a given data stream. One of the most fruitful of these relationships so far investigated for myocytes is the concept of cardiac restitution, the relationship between APD for a given beat to the diastolic interval (DI) of the preceding beat, DI = BCL−APD. Earlier work on this topic typically employ relationships of the form APD^*n*+1^ = *F*(DI^*n*^) which can be rewritten as APD^*n*+1^ = BCL *F*(APD^*n*^): = *G*(APD^*n*^) (Guevara et al., 1984; Kulkarni and Tolka-cheva, 2015; Kesmia et al., 2019). These approaches are heavily dependent on pacing protocols and a steady-state response with fixed BCL and as such have limited applicability in the case of spontaneously-beating tissue such as is observed in cultures of hiPSC-CMs. To investigate the behavior of cardiac restitution in spontaneous-beating cells with variable BCL using the language of discrete dynamical systems, we propose to consider

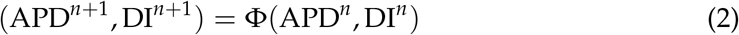

and observe these pairs from beat to beat, see Figure 8. Figures of this type suggest the presence of an underlying dynamical system for the two-dimensional state space. Further research on this topic may use deep neural networks to fit a function to these time series or exploit the field structure of the complex numbers to apply a convolutional neural network (Cantwell et al., 2019) to the number *z*^*n*^ = APD^*n*^ + *i* DI^*n*^.

**Figure 8:**
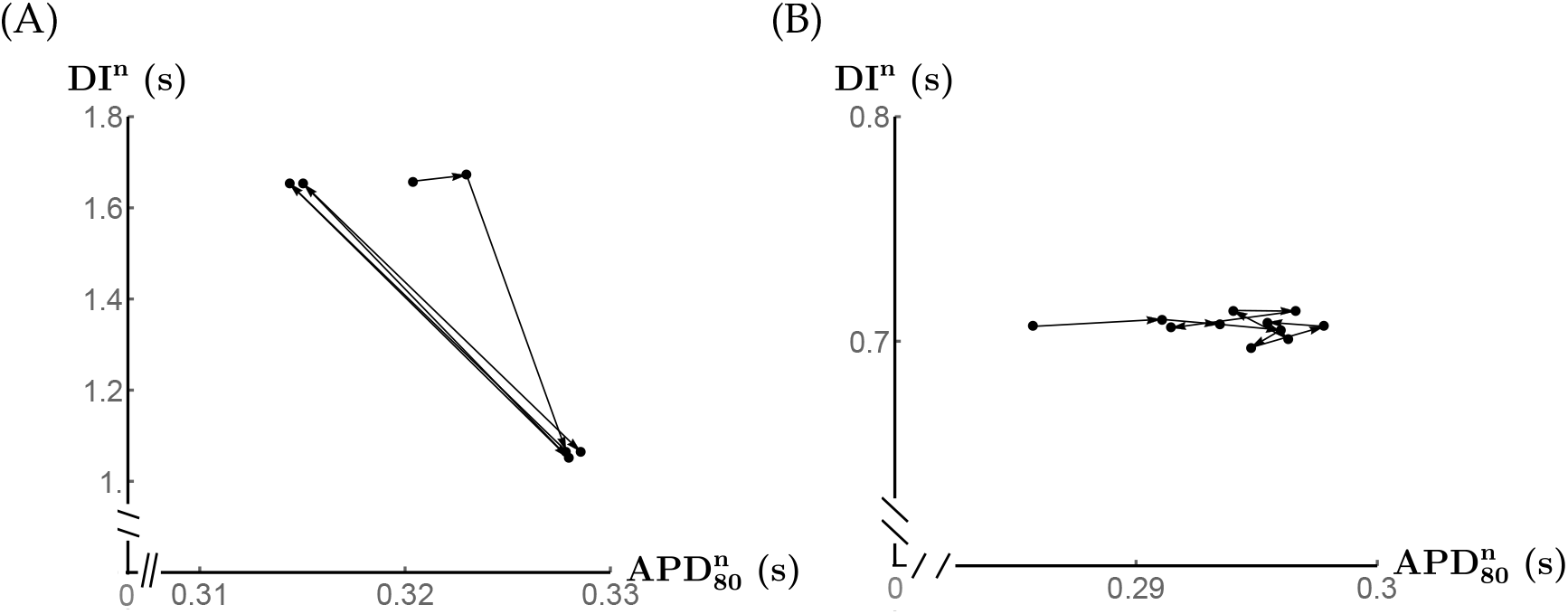
Dynamic plot of (APD_80_, DI) from two electrode streams for sequences of seven and 11 beats, respectively. Panel (A) shows alternans while (B) shows strong variability in the APD tied to a constant DI. Note that both point sets are largely confined to one-dimensional subspaces.

The above implementation only extracts a small number of numerical features primarily from AP traces for analysis. The segmentation algorithm creates data arrays that can be mined for any number of features, allowing user input as to which features to extract for a given application. In particular, FP duration is not computed by this program. This requires accurate identification of the repolarization wave in the FP complex. Automated detection of such waves is quite challenging, with several algorithms developed (Elgendi et al., 2015; Shang et al., 2019), but these come with computational overhead. MEA signals do not always manifest a recognizable repolarization wave. This may be tied to cell linkage to the substrate in the MEA (Li et al., 2017) or perhaps to immaturity of cells.

## Acknowledgments

We thank the Advocate Aurora Research Institute (Milwaukee, WI) for permission to use data that were generated there. We thank the editors and reviewers for valuable comments.

## Non-standard Abbreviations and Acronyms

AP: action potential
FP: field potential
MEA: multi-electrode array
hiPSC-CMs: human induced pluripotent stem cell derived cardiomyocytes
APD/APD_*m*_: action potential duration (to *m* % level)
BCL: basic cycle length
DI: diastolic interval
DFT: discrete Fourier transform
SVM: support vector machine

## Online Supplement

### *L*^2^ **and Supremum Norms**

Given a vector **x** = (*x*_1_,…, *x*_*N*_), the *L*^2^-norm of **x** is

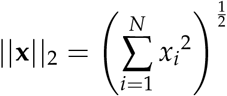

and the supremum, or *L*^∞^-norm

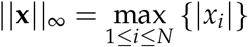

These norms define a distance function *d* between vectors by setting *d*⟨*x, y*⟩ = ||*x* − *y*||.

## Discrete Fourier Transform

Given a vector **x** = (*x*_1_,…, *x*_*N*_), we define its discrete Fourier transform by (Oppenheim and Schafer, 1999)

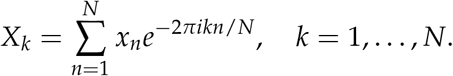

This is the discrete analog of the continuous Fourier transform, and the modulus |*X*_*k*_| indicates the relative contribution of sinusoids with frequency 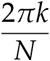 to the overall periodic signal.

## Supplementary Material

The light microscopy video (10X) “video.avi” shows spontaneously beating hiPSC-CMs at 24 days post-differentiation.

